# The Evidence of Cough Contagion in Human Beings

**DOI:** 10.1101/2023.05.25.542258

**Authors:** Ziyou Wang, Huaiqing Wang, Zihan Xia, Dazhi Gao

## Abstract

There is the evidence for cough contagion in human beings caused by empathy rather than physiology.Due to limited study on cough contagion,researchers have doubted whether cough was contagious from emtional contagion like yawn contagion.To deal with the doubts,we observed coughs from 34 adults in Ocean University of China in real time and recorded individual ‘s gender and local time.Then we developed a mathematical model to divide the cough process into several bouts and set a threshold for contagion to construct a response graph.With the graph,we first saw a strong effect of contagion for around 30 seconds no matter how long the bin(< 30*s*) was.Afterwards with mutiple measures,we extracted seven features(e.g.,duration) to describe the transmission chain and then found that there ‘s no time difference in cough contagion.Moreover,we also discovered tthe gender difference that males were more likely to be induced by triggers.Thus,cough contagion seems to be a normal phenomenon in human beings providing support to the experimental studies of empathy theory.

## 1. Introduction

Contagion is a rapid communication of an influence such as a behavior or emotional state in human beings. There are two main types of contagion,a physical one and an emtional one.Physical contagion,which is also called infection,usually develops by virus like recent COVID-19 and it ‘s increasingly linked to the development of the medical infrastructure and training ^[1]^. Alternatively,emtional contagion is often defined as behavioral and attentional synchrony resulting from social interactions ^[2]^. Past researches revealed that emotional contagion exists and causes mind and body arousal ^[3]^, which shows a similar facial, vocal,or postural expression,as well as similar neurophysiological and neurological reactions ^[4]^. The study of emotional contagion in human beings has awakened the interest of various research fields,such as human-robot interaction,decision-making in emergencies and commercial transactions ^[5, 6, 7, 8]^.

Emotional contagion theory is historically based on the theory of arousal ^[9, 10, 11]^,which reflects on the specific role in social interactions like triggers. Moreover,methodological approaches to emotional contagion are multiple,including facial expression reactions,behavioral reactions,physiological reactions and neurological reactions. Specifically,facial expression reactions ^[4, 12, 13, 14]^ are described as the one of the main methodological approaches used to study emotional contagion and behavioral reactions. Behavioral reactions come second,which has been proved that observing other people ‘s behavior,such as laughter ^[15, 16, 17]^,yawn ^[14, 15, 18]^and cough,on social networks triggers emotional contagion ^[19, 20]^.Generally,physiological and neurological reactions reveal that emtional contagion is associated with physiological reactions ^[21, 22]^, exactly the skin conductance response ^[4, 21, 22, 23]^. With the help of the neuroimaging tools like functional Magnetic Resonance Imaging (fMRI) ^[24]^, the root of emotional contagion were thought of being connected with various brain areas ^[25, 26, 27]^. Adopting the method of facial expression and behavioral reactions,emtional contagion is divided into something more concrete. For instance,many researchers have underlined the salience of yawn contagion among human beings,which has been proved existent in many experiments ^[28, 29, 30]^.

Contagious yawning is when one individual ‘s(termed here: the trigger) yawning induces a yawn in another one(termed here: the obeserved) who would not have yawned ^[31]^. Evidence for the existence of yawn contagion and other by-conclusions in humans have historically originated from the analysis of observational data from a statistical perspective. What ‘s more, yawn contagion is essentially observed by facial expression reactions,which is more of vision contagion out of others like audition. According to this,past researchers drawed various conclusions on yawn contagion. In biological evolution,yawn contagion has indicated that emotional affinity matters more than speices by comparing humans (*Homo sapiens*) and bonobos(*Pan paniscus*) [18];In psychobiology,the occurence of yawn contagion have been obeserved in preschool children ^[32]^;In behavior and physiology,a critique and quantification of yawn contagion has been given ^[33]^. Besides the mainstream observation on visual yawn contagion,there are also some literatures ^[14, 30]^ about auditory yawn contagion,but it turns out that vision is more contagious than audition. Generally,most of the studies of yawn contagion relies on the aim of studying social bonds in groups ^[14]^, children development ^[34, 32]^, the mirror neuron system(MNS) ^[35, 36]^ and other interdiscipline.

Yawn contagion research has its own advatanges: representativeness,operability and scalability. That ‘s why researches on certain behavioral contagion are followed by the yawn contagion,which can also be described as an exmaple when dealing with emtional contagion issues. Although yawn contagion research stands out,there are still some difficulties to be conquered. The most obvious one is that yawn contagion research was limited to visual observation,which is hard to judge whether the potential observers get the stimuli from the triggers. To avoid this controversy and carry on their studies,yawn contagion researchers typically define a similarly ‘Distant view ‘ ^[31]^, which within it observers were assured to get induced by triggers. But when it comes to auditory observation,there is no need to worry about the ‘Distant view ‘ as sounds can be noticed at every corner of a relatively small room with reverberation. Correspondingly, especially in the post-pandemic era,cough,mainly observed by audition,has been considered to be normal physiological response just like yawn. The question whether cough is contagious arouses great interest.

Cough(or the cough reflex) is the defensive reflexes from the respiratory tract,which are elicited by mechanical and chemical irritation of the airway mucosa ^[37]^. Cough has been traditionally considered as a infectious disease and the cough contagion was partly thought as a physical contagion ^[38, 39]^ while possibility that cough contagion are related to social interactions and psychological factors ^[40, 41]^ was first proposed,which means cough contagion can be emtional. Representatively,experiments on college students preliminarily concluded that people were more likely to cough if they heard others cough ^[40]^. However,former study just compared the mean of coughs per minute with the expected constant cough frequency. Without describing the contagion process mathematically,the conclusion was not rigorous enough,which in some degree left some space for following study. Even though,nowadays cough contagion research are not as popular as other researches on similar categories. The main reason is that cough is much more easily to be inhibited or abolished ^[42]^ than other spontaneous physiological activities,which makes it hard to observe cough events. Furthermore,considering the social distance ^[43, 44]^ cough brings,observational data in public,especially among strangers,are hard to obtain.

This paper focuses on finding the evidence of cough contagion in human beings using a mathematical model. Specifically,based on the findings from Pennebaker ^[40]^ and contagion judgment model similar to yawn contagion from Campbell ^[31]^, the model proposed by this paper analyzed and quantified the whole contagion process,such as trigger, observer, time interval, then defined the criterion whether this process reflected contagion. Additionally,time factors(moring or afternoon),difficult to analyze though,were also taken into consideration. In this study,we hope to supplement and expand the theory of emotional contagion,reveal that cough contagion is a natural phenomenon in human society,and provide data support for subsequent experiments and observations in discipline such as behavioral science and cognitive neuroscience.

## 2. Method

### 2.1. Ethic

All participants voluntarily participated in the experiment and signed an informed consent form before the experimentand could receive a certain amount of compensation after the experiment. The data collection protocol was approved by Ocean University of China.

### 2.2. Assumptions

For the convenience of observational data collection and analysis,we made one assumption that due to the limited number of experimental personnel,limited venue space,and strong reverberation,cough will be heard or noticed by all other participants.

### 2.3. Subjects

We studied 34 students (17 females and 17 males,age range: 21–22 years old) who come from the same class and have social interactions with each other for around 30 months. All of the students are individuals with no serious physical abnormalities,and are in good mental state. The enviroment was the classroom where they were attending one specialized course. One thing needed to be emphasized is that not everyone was present in each observation.

### 2.4. Obeservations

All observations took place between two periods of time within a day which are 8:00-10:00 and 13:30-15:30. Researchers were separately located in different corners to make sure everyone in the room can be recorded. Data were taken by both monitor and handwritings of researchers in time series.

Observations include all coughs that occur in real time. Data was collected in real time by researchers and checked against each other to avoid omissions and errors. When a cough event occured,the researcher recorded the time of cough occurrence (accurate to seconds) and the corresponding individual(position encoded). According to all these data observation rules,researchers respectively collected the cough data.

### 2.5. Operational definitions and criterion

Given that cough contagion process hasn ‘t been quanitified,we gave several definitions,which were similar to those in yawn contagion,to describe cough contagion process. Cough often occured in bouts^[45]^, which sequentially consisted of Trigger Cough(TC),Middle Product of Contagion(MPOC) and Ending Cough(EC). Different from yawn contagion,we defined effective duration of self contagion(**termed here:***T*_1_) and mutual contagion(**termed here:***T*_2_) to accurately describe TC,MPOC and EC.

In order to describe the model,the occurrence time of a certain cough was defined as *t*. Here ‘s the detailed explanations of given definitions and criterions.

#### 2.5.1. Duration of Self and Mutual Contagion

*T*_1_ defined one type of the cough that was induced by oneself. This definition was innovative as we discovered that one individual could produce continuous cough out of others ‘ influence. We supposed it unviseral and measured the intervals between continuous coughs from the same individual. *T*_2_ shared the similar definition. The only difference was that measured intervals were the gap between continuous coughs from the closest individual in time. According to the data,we took the duration with the highest frequency of occurrence as the values of *T*_1_ and *T*_2_ respectively,where *T*_1_ was 30 seconds and *T*_2_ was 45 seconds.

#### 2.5.2. Lonely Cough

During the contagion,abnormal coughs were those arose independently,which can be defined as ‘Lonely Cough ‘. When someone coughed and there ‘s no mutual cough within (*t* − *T*_2_, *t* + *T*_2_),and there ‘s no self cough within (*t* − *T*_1_, *t* + *T*_1_),the cough was a lonely cough,which needed to be dropped. There were two exceptional cases while judging the lonely coughs.

- When the cough was the first one during the process,if there were no other mutal coughs after *T*_2_ and no self coughs after *T*_1_,this cough could be defined as lonely cough.
- When the cough was the last one during the process,if there were no other mutal coughs before *T*_2_ and no self coughs before *T*_1_,this cough could be defined as lonely cough.

#### 2.5.3. TC, EC and MPOC

There were two main methods to decide *TC*. On one hand, *TC* was the first cough in the process. On the other hand,when there were no mutual coughed within (*t* − *T*_2_, *t*) and no self coughed within (*t* − *T*_1_, *t*),the cough could also be defined as *TC*.

There were also two main methods to decide *EC*. Firstly, *EC* was the last cough in the process. Secondly,when there were no mutual cough within (*t, t*+*T*_2_) and no self cough within (*t, t*+*T*_1_),the cough could also be defined as *EC*.

Cough often occured in bouts^[45]^, which sequentially consisted of Trigger Cough(TC),Middle Product of Contagion(MPOC) and Ending Cough(EC). Trigger Cough served as the starting point of the bout,Ending Cough served as the ending point,and Middle Products of Contagion were between them.

#### 2.5.4. Process for Special Scenario

Although we had divided the cough contagion process into bouts,there were also some scenarios in the observation. When it came to the situation that the same individual had two continuous coughs with a time difference less than *T*_1_ in the same bout,we could neither tell which of the two cough induced the following cough from others nor tell whether the second cough was induced by others(mutual contagion) or the first cough(self contagion). Those problems would bring about some exceptional cases.

To deal with exceptional cases without influencing the original features(contagion) of obervational data,we chose to use the most conservative approach,which decreased MPOCs as many as possible by transforming suitable MPOCs into *TC* or *EC*. We named it Process for Special Scenario(PSS). Next,we would elaborate on different exceptional cases and corresponding solutions.

##### Case I

After one individual gave *TC*,we couldn ‘t tell whether the first MPOC coming after the *TC* from the same individual was the real MPOC when it was within (*t, t* + *T*_1_).

To deal with it,we proposed two methods. One method was that when there was no others ‘ coughs between *TC* and false MPOC,we dropped the original *TC* and replaced it with the false MPOC. The other method was that when there were others ‘ coughs between *TC* and false MPOC,we would transform the cough of others which was closest to the false MPOC in time into *EC* of this bout and change the false MPOC into *TC* of the following bout.

##### Case II

After one individual gave MPOC,we couldn ‘t tell whether the first MPOC coming after the MPOC from the same individual was the real MPOC when it was within (*t, t* + *T*_1_).

To deal with it,we proposed two methods. One method was that when there was no others ‘ coughs between the first MPOC and the second MPOC,we would transform the first MPOC into the *EC* of this bout and the second MPOC into the *TC* of the following bout. The other method was that when there were others ‘ coughs between the first MPOC and the second MPOC,we would transform the cough of others which was closest to the second MPOC in time into *EC* of this bout and the second MPOC into *TC* of the following bout.

##### Case III

After one individual gave MPOC,we couldn ‘t tell whether the *EC* coming after the MPOC from the same individual was the real *EC* when it was within (*t, t* + *T*_2_).

To deal with it,we proposed two methods. One method was that when there was no others ‘ coughs between the false MPOC and *EC*,we would drop the original *EC* and replace it with the false MPOC. The other method was that when there were others ‘ coughs between the false MPOC and *EC*,we would transform the cough of others which was closest to the MPOC into *TC* of this bout and the false MPOC into *EC* of the previous bout.

### 2.6. Procedure

Before analyzing the data to test and verify the cough contagion,we needed to preprocess the original observational data. In the beginning,we dropped coughs from two or more individuals at the same time and then filtered out lonely coughs. Next,we decided the TC,EC and MPOC. After this,we took the exceptional cases into considertion and then got the initial bouts,including each TC,EC and MPOC. Subsequently,we determined whether Process for Special Scenario(PSS) was required in chronological order in the time series. After each PSS,we should observe the new data to reassess whether we needed to perform PSS again. That ‘s the end of preprocessing.

Having obtained the qualified data,we divided the whole into numerous bouts based on TC and EC and then perfromed the stats in each bout. We supposed 5 seconds as a time bin for statistic analysis. This number is arbitrary,as we had no data from any researches.

In one bout,we moved the bin from TC to EC in order and counted the number of MPOCs and ECs in each bin that belonged to the bout in sequence. That ‘s to say,we did not count all types of cough in the last bin of this bout in the next bout,as well as the TCs. Analyzing all the bouts separately in the same way we proposed and then adding the number of MPOCs and Ecs counted in different bouts at the same time point together,we changed the messy observational data into a 1 × *N* numerical matrix,

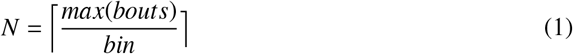

where,*max*(*bouts*) is the longest duration among all the bouts.

Afterwards,we calculated the mean value *µ* and standard deviation σ of cough number in bouts and we set a threshold to allow 4.95% of bin to exceed,supposing the data conformed to the Z distribution. At last when one certain exceeded the threshold,we considered that such bin showed contagion. Up to now,we thought that our criteria were the most conservative method to judging whether cough contagion could be detected in observational data. In addition,while judging the contagion,we had set false positive and false negative controls in the results to ensure their reasonableness. When the first few bins were all considered contagious,the process of cough showed contagion. Other words,if the contagious bin did not appear continuously in the beginning,there would be no contagious cough.

#### False positive control

Because we calculated so many bins,there was a possibility that a bin could be above threshold simply due to the calculating error. If a bin,which was isolated from others,was above threshold,it would not make sense to conclude that coughs were contagious in that bin. Thus,we formed a priori rule,in regard to false negative,that when a bin exceeded the threshold,but neither of its surrounding bins exceeded the threshold,it was considered that the bin was not contagious.

#### False negative control

There was also the possibility for the reverse. In the same way,when a bin didn ‘t exceed the threshold,but both its surrounding bins exceeded the threshold,it was considered contagious.

## 3. Results

### 3.1. Evidence of cough contagion

We observed students for a total of 400 minutes across 8 sessions. In this time,we recorded 390 coughs at an overall rate of 0.98 coughs per minute. These coughs finally occured in 82 bouts with an average of 2.85 coughs(SD=1.20).

To describe whether the coughs by bouts implied contagion,we set 5 seconds as a bin and then counted the total number of coughs in different bouts at the same time point(See Figure.1). From the histgram,we could know that number of coughs had exceeded the threshold,which initially reflected the contagion. Although the whole process had been divided into several mutually independent bouts,each bout demonstrated similar conclusion that only the first six bins were contagious,which showed the process of coughing was contagious and contagion lasted for 30 seconds.

**Figure 1:**
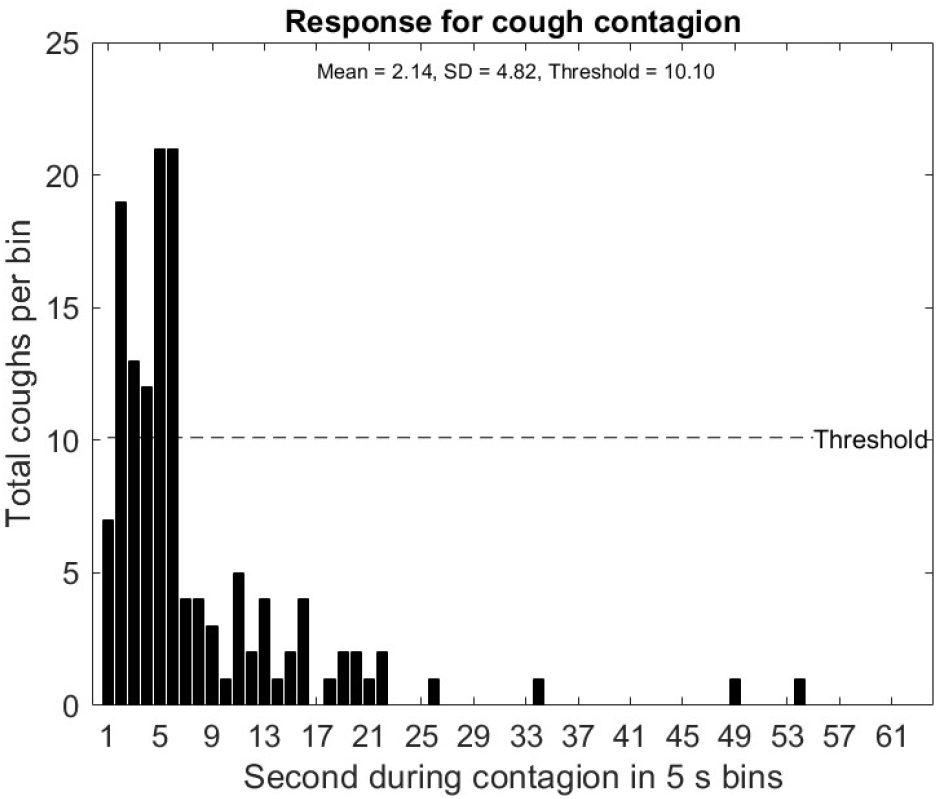
A graph of total number of coughs when bin was 5 seconds in each bout. PSS had been applied to ensure the most conservative result of cough contagion. The mean was 2.14 per bin with a standrad deviation of 4.82. To calculate the threshold we took the mean and added 1.65 times the standard deviation,yielding a threshold of 10.10 coughs per bin(p < 0.05 on a z distribution). With the help of threshold and our criteria for false positives and negatives,we saw evidence for cough conatgion within 30 s.

To make sure the bin we set would not change the general result that the cough contagion was existent,we adjusted the time window(bin) to 3s(Fig.2(a)),10s(Fig.2(b)),15s(Fig.2(c)) and 20s(Fig.2(d)) and found out the results still had strong effect(Figure.2),all lasting for around 30s.

**Figure 2:**
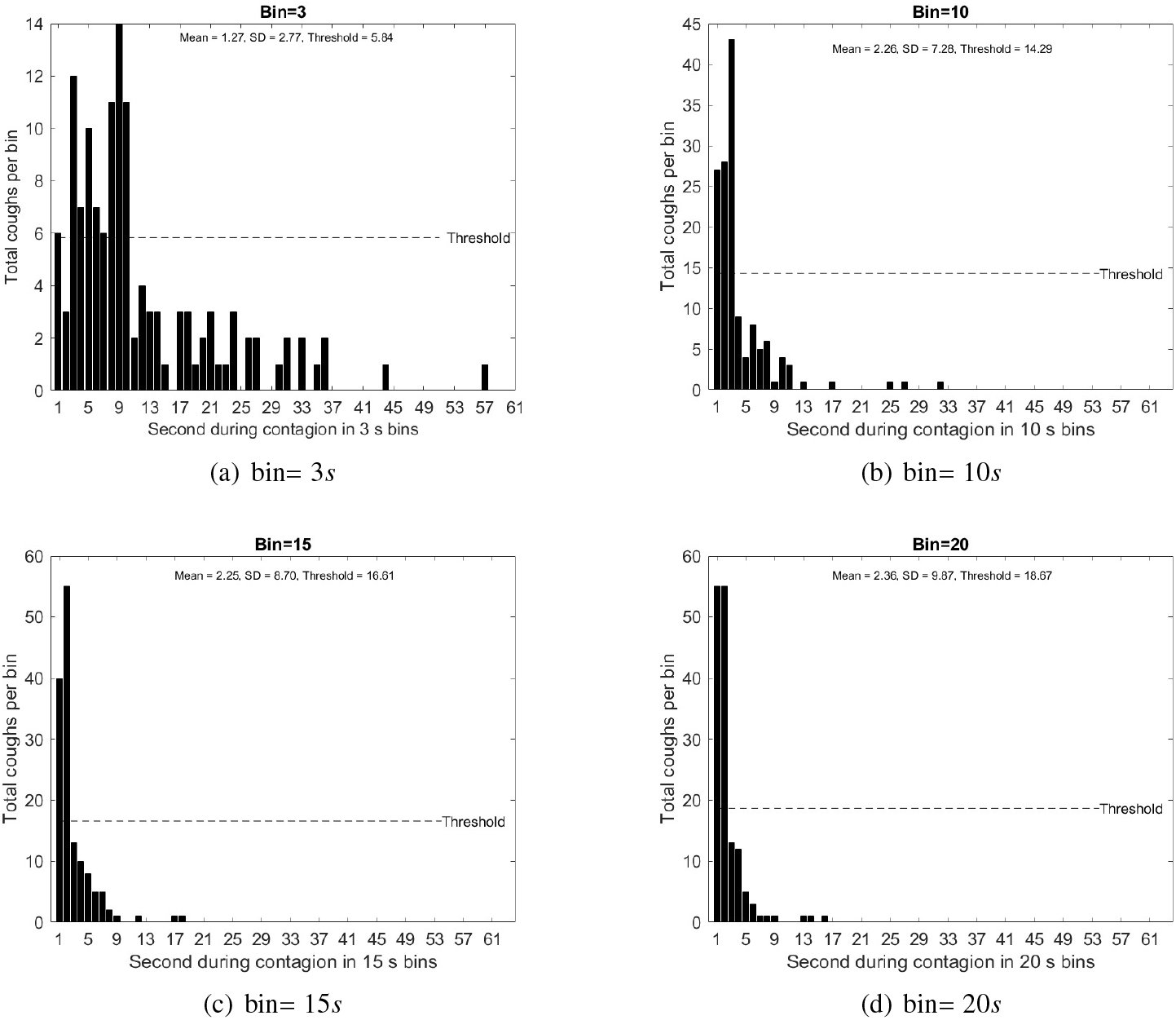
**(a-d)** After changing the value of bin to 3*s*,10*s*,15*s* and 20*s*,the patterns were similar to the one in 5*s*,which showed a strong effect in cough contagion and the durations were always around 30*s*.

### 3.2. Chain of transmission

PSS was a tool to simplify the judgement of cough contagion. When it came to the properties of contagious coughs,we needed to count the component of each bout,which meant we needed to analyze the data before PSS. Since we had divided the contagion into several bouts,in which were the sets of proved contagious coughs(See 2.5.4),those bouts could be also viewed as chains of transmission. To better describe the chains,we extracted five numerical features based on the observational features(time,cougher number):**cough amount**(mean = 2.85,SD = 1.20) was defined as the amount of coughs in each chain,which represented the length of each chain;**response time**(mean = 15.43,SD = 8.07) was defined as time cell(seconds) between first two coughs,which represented the transmission rate;**duration**(mean = 44.00,SD = 55.69) was defined as the time cell(seconds) between the first cough and the last cough,which represented the action time of each chain;**diversity**(mean = 2.39,SD = 0.77) was defined as the types of cougher,which represented the richness of one chain;**density**(mean = 0.23,SD = 0.08) was defined as the ratio of diversity to total cough numbers,which represented the reach of each chain.

We used Correlation Analysis to observe the internal connections of the transmission chain in order to indicate the features of cough contagion. Since the data were not tested normally distributed,we used nonparametric correlation. There was a significant correlation between the cough amount and duration(Spearman ‘s rho = 0.792, p < 0.01),meaning that the cough amount increased as the duration accumulated. There was also a significant and positive correlation between response time and duration(Spearman ‘s rho = 0.339, p = 0.002 < 0.01),which revealed that the longer response time was,the longer the cough contagion would last for. Besides the duration showed importance in cough contagion,the diversity was both significantly correlated with cough amount(Spearman ‘s rho = 0.747, p < 0.01) and duration(Spearman ‘s rho = 0.608, p < 0.01),indicating that more different individuals engaged in the cough contagion,the longer the transmission chains would last for. What ‘s even more interesting was that density might be the best feature to describe transmission chains as it was highly positively correlated with cough amount(Spearman ‘s rho = 0.339, p < 0.01),response time(Spearman ‘s rho = 0.222, p = 0.045 < 0.05),duration(Spearman ‘s rho = 0.520, p < 0.01),diversity(Spearman ‘s rho = 0.716, p < 0.01).Except the obvious conclusion that diversity was connected with density,the large density brought about more coughs,longer response time and longer duration in one chain. Nonetheless,we could further examine cough contagion effects in the following discussion.

### 3.3. Time difference of contagion

We observed two groups of data from the same individuals at different times(see Method),which divided the 82 bouts into 48 bouts in A.M(mean = 2.90,SD = 1.26) and 34 bouts in P.M(mean = 2.79,SD = 1.12). To explore time difference of cough contagion and where the differences located,we used One-way Analysis of Variance(ANOVA) to analyze each feature,which were raised above. We wondered whether these five features had differences on time scale. In detail,we first analyzed if there was a significant difference between time(A.M,P.M) and features(cough amount,response time,duration,diversity and density). Then we described the specific differences by comparing the average values in detail.

Based on ANOVA,there was no significant difference between cough number in A.M and P.M(mean A.M = 2.79, mean P.M = 2.90,F = 0.142,p = 0.707 > 0.05). So were the response time(mean A.M = 14.88, mean P.M = 15.81,F = 0.262,p = 0.610 > 0.05),duration(mean A.M = 36.03, mean P.M = 49.65,F = 1.193,p = 0.278 > 0.05),diversity(mean A.M = 2.24, mean P.M = 2.50,F = 2.419,p = 0.124 > 0.05) and density(mean A.M = 0.23, mean P.M = 0.23,F = 0.040,p = 0.843 > 0.05). As a whole,all raised features showed no significant difference on a time level,which also implied that the process of cough contagion was independent of time changes.

### 3.4. Sex difference of contagion

We also observed the individual gender and tried to tell the sex difference in cough contagion. One thing that needed to be emphasized was that we divided the total bouts into male cough(mean = 3.02,SD = 1.39) and female cough(mean = 2.65,SD = 0.89) bouts according the trigger ‘s gender in each bout. We also used ANOVA to research whether the five features had differences on sex scale. In order to find out more sex difference,we added to two more features,which were **homosexual contagion number** and **heterosexual contagion number**. Both of them showed the characteristics(heterosexual or homosexual) of gaps of one chain,and we counted separately the total number of the gap,which represented the transmission from the same or different gender,from the adjacent coughs as homosexual contagion number and heterosexual contagion number. For example,if one chain contains six coughs(five gaps) with its sexual series 100001(male in 0 while female in 1),there would be three homosexual contagion and two heterosexual contagion.

Gathering all seven features,we adopted ANOVA as what we did in the research of time difference. There were no significant difference between cough number in males and females(mean male contagion = 3.02, mean female contagion = 2.65,F = 1.99,p = 0.16 > 0.05),response time(mean male contagion = 15.42, mean female contagion = 15.43,F = 0.00,p = 0.995 > 0.05),duration(mean male contagion = 54.33, mean female contagion = 31.43,F = 3.54,p = 0.064 > 0.05),diversity(mean male contagion = 2.40, mean female contagion = 2.38,F = 0.016,p = 0.900 > 0.05) and density(mean male contagion = 0.24, mean female contagion = 0.22,F = 0.407,p = 0.525 > 0.05). Those data showed that five basic features,which contained information of transmission chains,had nothing to do with sex difference. After adding the new features,we found that there was significant differences between homosexual contagion number in males and homosexual contagion number in females(mean male contagion = 0.73, mean female contagion = 0.48,F = 7.829,p = 0.006 < 0.01). So was heterosexual contagion number(mean male contagion = 0.27, mean female contagion = 0.49,F = 6.208,p = 0.015 < 0.05). Having compared the average value of homosexual contagion number and heterosexual contagion number,we concluded that contagious cough induced by males were more likely to cause homosexual contagion while contagious cough induced by females were more likely to cause heterosexual contagion.The specific reasons will be discussed below.

## 4. Discussion

The results show evidence that cough contagion exists in human beings and the effect is strong(the peak of the graph> 5 standard deviation).When we adjusted the value of bin under 30s,we could get the same conclusion that the contagion still was existent and lasted for 30 seconds(Figure.2). Although we have drawn the conclusion,researches on the cough contagion are limited compared to other contagion of facial expression ^[46]^ such as yawn and laughter ^[15]^. The only study on cough contagion was conducted by Pennebaker ^[40]^, who used ordinary chi square test to support the observation of cough contagion. Our study can be a great supplement to his research by analyzing the cough at a finer level,transmission chains actually. Since the cough contagion has been proved existence,we can respond to doubts whether cough can be infected by social factors. We may also support the hypothesis by Platek ^[47]^, and extend to the conclusion contagious cough is a byproduct of evolutionarily designed programs related to empathy, social cognition and self-awareness. We possibly regard cough contagion as embody of empathic processing. In an fMRI study ^[46]^, empathic processing(mainly yawn contagion) is associated with ventromedial prefrontal cortex (vmPFC) activation,which has been indirectly shown to be the human mirror neuron system ^[48]^. From the behavioral data in our study,we propose a conjecture that cough contagion is similar to yawn at the level of brain area. Due to limitations in experimental conditions,we cannot confirm our conjecture without functional MRI data.

After we proved the existence of cough contagion,we wonder the characteristic of it or how it will act on human society. Thus,we focus on the process of cough contagion,which is transmission chain or we can equate it to the bout in our study. We extracted five basic features(cough amount,response time,duration,diversity and density) and two extended features(homosexual contagion number and heterosexual contagion number)from observational data. We highlighted the density as the most significant role in contagion because it is significantly and positively correlated with other four basic features,which shows that with more individuals engaged in the contagion process,the transmission chain will be longer in time,richer in member composition and more in containing coughs. That ‘s an obvious conclusion as it has been mentioned before ^[40]^. In addition,we make one interesting discovery that longer response time might brings more individuals induced by trigger. We possibly attribute this discovery to mirror neuron system,which has been proved related to somatosensory, auditory and emotional processing ^[49]^. When the response time tends to be longer,which means more time for mirror neurons to perceive the action ^[50]^, more observing individuals will recognize and understand it,leading to multiple responses as a result. Rigorously,more fMRI and EEG evidences are required to support our explanation.

As contagion is highly connected with two social factors(environment and observers),we try to use the features above to tell the difference in cough contagion. We choose time(A.M and P.M) on behalf of enviroment and gender(male and female) on behalf of individuals despite that they may not fully reflect differences,but we suppose they have wider applicability,which can better describe the cough contagion in emtional contagion theory ^[9, 10, 11]^.

Traversing all features above,we draw the conclusion that the cough contagion,transmission chains set to be exact,have nothing to do with time(all *p*>0.05 in ANOVA). Although cough itself has time perferrence,which means that coughs are more likely to occur in the midafternoon ^[51]^, cough contagion shows no difference in time scale. To some extent,it proves again that cough contagion is more emtional than physical.

Then we turn to individuals ‘ gender to tell whether there are differences in sex and we honestly find something impressive. Among seven listing features,whether the contagion appears between the same sex or not really shows significant difference between males and females. For homosexual contagion number,the difference is significant and the males take the upper hand(mean male contagion = 0.73, mean female contagion = 0.48,F = 7.829,p = 0.006 < 0.01). For heterosexual contagion number,the difference is significant and the females win(mean male contagion = 0.27, mean female contagion = 0.49,F = 6.208,p = 0.015 < 0.05). From these,we can conclude that when the trigger is male,more homosexual contagions will appear while when the trigger is female,more heterosexual contagion will occur. In one word,males are much easily to be infecetd than females. However,gender differences in mirror neuron system specifically reveals that affective arousal and expression of emotion (e.g., in response to the emotions of other people) demonstrate superior performance of females over males ^[52]^. This contradiction may be caused by the female psychology,which implies even if females have the willing to cough,they first consider the social distance the coughs bring ^[43]^ and then voluntarily control themselves ^[42]^ to avoid the occurence of coughs. It is one potential explanation and there is another possibility that our data volume is too small to significantly highlight gender differences.

To summarize,these conclusions we drew are direct responses to the mathematical model we have established. Although we have gotten some preliminary results,such as study on transmission chain,analysis of time difference and sex difference,there is still much to improve. First of all,more valid data with large volume and high dimension like attentive level are required. Secondly,the evaluation of model parameters need to be quantified,which means we should give more evidence to explain why the parameter like bin takes this value. Thirdly,to make our new discoveries like sex difference more accurate,we should connect our behavior results calculated by the model with neuroimaging results such as fMRI,EEG and MEG as the cough contagion essentially the neural activity of the human brain.

We hope our study can bring inspiration to more contagion study. From a model perspective,we provided a new thought to handle auditory information and a relatively reasonale model to determine contagion which is not limited to cough. From the perspective of conclusion analysis,we closely combined emotional contagion with the mirror neuron system,which possibly provides the neurological basis of human self-awareness ^[53]^.

## 5. Conclusion

Based on the observing cough data we collected,we gave some definitions to describe the process of continous cough incidents and one criterion to determine whether cough process was contagious. Having drawn the conclusion that cough contagion is a natural phenomenon in human beings and it is possibly reflection of mirror neuron system,we extrated some features(cough amount,response time,duration,diversity density,homosexual contagion number and heterosexual contagion number)from the transmission chains to further analyze the characteristics of contagion. Correlation analysis were used to conclude that density was the most significant feature to describe contagion and longer response time increases density due to action understanding theory. Then ANOVA were used to tell the time and sex difference in cough contagion,which demonstrated that there was no time difference in spite of cough itself having time perferrence. Moreover,gender difference appeared in the mode of transmission(homosexual or heterosexual),which showed that males were much easier to be influenced than females.Considering the small data volume,low data dimension and single scientific method without neuroimaging,our study might not handle the details well but it provided general conclusions about cough contagion.

## Acknowledgments

We would like to acknowledge the assistance of volunteers in putting together this example manuscript and supplement.

## Author Contributions

Z.W,H.W and D.G designed the study. Z.W,H.W and Z.X collected data. Z.W and H.W analyzed the data and wrote the paper. Z.X designed algorithm.

