## Supplementary material for "The Evidence of Cough Contagion in Human Beings": Code: account.m

```
clc;
clear;
% close all;
sum_N1=load("sum_N_3.19_1.mat");
sum_N2=load("sum_N_4.14_1.mat");
sum_N3=load("sum_N_4.14_2.mat");
sum_N4=load("sum_N_4.18_1.mat");
sum_N5=load("sum_N_4.18_2.mat");
sum_N6=load("sum_N_4.23_1.mat");
sum_N7=load("sum_N_4.23_2.mat");
sum_N8=load("sum_N_4.25_1.mat");
sum_N9=load("sum_N_4.25_2.mat");


sum_N2_sex_0=load("2_sex0.mat");
sum_N3_sex_0=load("3_sex0.mat");
sum_N4_sex_0=load("4_sex0.mat");
sum_N5_sex_0=load("5_sex0.mat");
sum_N6_sex_0=load("6_sex0.mat");
sum_N7_sex_0=load("7_sex0.mat");
sum_N8_sex_0=load("8_sex0.mat");
sum_N9_sex_0=load("9_sex0.mat");

sum_N2_sex_1=load("2_sex1.mat");
sum_N3_sex_1=load("3_sex1.mat");
sum_N4_sex_1=load("4_sex1.mat");
sum_N5_sex_1=load("5_sex1.mat");
sum_N6_sex_1=load("6_sex1.mat");
sum_N7_sex_1=load("7_sex1.mat");
sum_N8_sex_1=load("8_sex1.mat");
sum_N9_sex_1=load("9_sex1.mat");

sum_N1=sum_N1.sum_N;
sum_N2=sum_N2.sum_N;
sum_N3=sum_N3.sum_N;
sum_N4=sum_N4.sum_N;
sum_N5=sum_N5.sum_N;
sum_N6=sum_N6.sum_N;
sum_N7=sum_N7.sum_N;
sum_N8=sum_N8.sum_N;
sum_N9=sum_N9.sum_N;

sum_N2_sex_0=sum_N2_sex_0.sum_N;
sum_N3_sex_0=sum_N3_sex_0.sum_N;
sum_N4_sex_0=sum_N4_sex_0.sum_N;
sum_N5_sex_0=sum_N5_sex_0.sum_N;
sum_N6_sex_0=sum_N6_sex_0.sum_N;
sum_N7_sex_0=sum_N7_sex_0.sum_N;
sum_N8_sex_0=sum_N8_sex_0.sum_N;
sum_N9_sex_0=sum_N9_sex_0.sum_N;

sum_N2_sex_1=sum_N2_sex_1.sum_N;
sum_N3_sex_1=sum_N3_sex_1.sum_N;
sum_N4_sex_1=sum_N4_sex_1.sum_N;
sum_N5_sex_1=sum_N5_sex_1.sum_N;
sum_N6_sex_1=sum_N6_sex_1.sum_N;
sum_N7_sex_1=sum_N7_sex_1.sum_N;
sum_N8_sex_1=sum_N8_sex_1.sum_N;
sum_N9_sex_1=sum_N9_sex_1.sum_N;

[~,x1]=size(sum_N1);
[~,x2]=size(sum_N2);
[~,x3]=size(sum_N3);
[~,x4]=size(sum_N4);
[~,x5]=size(sum_N5);
[~,x6]=size(sum_N6);
[~,x7]=size(sum_N7);
[~,x8]=size(sum_N8);
[~,x9]=size(sum_N9);

[~,x2_sex_0]=size(sum_N2_sex_0);
[~,x3_sex_0]=size(sum_N3_sex_0);
[~,x4_sex_0]=size(sum_N4_sex_0);
[~,x5_sex_0]=size(sum_N5_sex_0);
[~,x6_sex_0]=size(sum_N6_sex_0);
[~,x7_sex_0]=size(sum_N7_sex_0);
[~,x8_sex_0]=size(sum_N8_sex_0);
[~,x9_sex_0]=size(sum_N9_sex_0);

[~,x2_sex_1]=size(sum_N2_sex_1);
[~,x3_sex_1]=size(sum_N3_sex_1);
[~,x4_sex_1]=size(sum_N4_sex_1);
[~,x5_sex_1]=size(sum_N5_sex_1);
[~,x6_sex_1]=size(sum_N6_sex_1);
[~,x7_sex_1]=size(sum_N7_sex_1);
[~,x8_sex_1]=size(sum_N8_sex_1);
[~,x9_sex_1]=size(sum_N9_sex_1);


max_num=max([x1,x2,x3,x4,x5,x6,x7,x8,x9,...
    x2_sex_0,x3_sex_0,x4_sex_0,x5_sex_0,x6_sex_0,x7_sex_0,x8_sex_0,x9_sex_0,...
    x2_sex_1,x3_sex_1,x4_sex_1,x5_sex_1,x6_sex_1,x7_sex_1,x8_sex_1,x9_sex_1]);
sum_all=zeros(1,max_num);

sum_all(1:x2)=sum_N2+sum_all(1:x2);

sum_all(1:x3)=sum_N3+sum_all(1:x3);

sum_all(1:x6)=sum_N6+sum_all(1:x6);

sum_all(1:x7)=sum_N7+sum_all(1:x7);

sum_all(1:x4)=sum_N4+sum_all(1:x4);

sum_all(1:x5)=sum_N5+sum_all(1:x5);

sum_all(1:x8)=sum_N8+sum_all(1:x8);

sum_all(1:x9)=sum_N9+sum_all(1:x9);

%男
% sum_all(1:x2_sex_0)=sum_N2_sex_0+sum_all(1:x2_sex_0);
% 
% sum_all(1:x3_sex_0)=sum_N3_sex_0+sum_all(1:x3_sex_0);
% 
% sum_all(1:x4_sex_0)=sum_N4_sex_0+sum_all(1:x4_sex_0);
% 
% sum_all(1:x5_sex_0)=sum_N5_sex_0+sum_all(1:x5_sex_0);
% 
% sum_all(1:x6_sex_0)=sum_N6_sex_0+sum_all(1:x6_sex_0);
% 
% sum_all(1:x7_sex_0)=sum_N7_sex_0+sum_all(1:x7_sex_0);
% 
% sum_all(1:x8_sex_0)=sum_N8_sex_0+sum_all(1:x8_sex_0);
% 
% sum_all(1:x9_sex_0)=sum_N9_sex_0+sum_all(1:x9_sex_0);

%女
% sum_all(1:x2_sex_1)=sum_N2_sex_1+sum_all(1:x2_sex_1);
% 
% sum_all(1:x3_sex_1)=sum_N3_sex_1+sum_all(1:x3_sex_1);
% 
% sum_all(1:x4_sex_1)=sum_N4_sex_1+sum_all(1:x4_sex_1);
% 
% sum_all(1:x5_sex_1)=sum_N5_sex_1+sum_all(1:x5_sex_1);
% 
% sum_all(1:x6_sex_1)=sum_N6_sex_1+sum_all(1:x6_sex_1);
% 
% sum_all(1:x7_sex_1)=sum_N7_sex_1+sum_all(1:x7_sex_1);
% 
% sum_all(1:x8_sex_1)=sum_N8_sex_1+sum_all(1:x8_sex_1);
% 
% sum_all(1:x9_sex_1)=sum_N9_sex_1+sum_all(1:x9_sex_1);

mean_N=mean(sum_all);

std_N=std(sum_all);

threshold_all=mean_N+1.65*std_N;

ture_infect=zeros(1,max_num);
ture_infect(find(sum_all>=threshold_all))=sum_all(find(sum_all>=threshold_all));
modle_judge=zeros(1,max_num);
modle_judge_res=zeros(1,max_num);
modle_judge(find(sum_all>threshold_all))=1;
%False negative judgment
for n=2:1:max_num-1
    point_front=modle_judge(n-1);
    point=modle_judge(n);
    point_back=modle_judge(n+1);
    if point_front==1 && point_back==1 && point==0
        modle_judge_res(n)=1;
        modle_judge(n)=1;
    end
end

%False positive judgment
for n=2:1:max_num-1
    point_front=modle_judge(n-1);
    point=modle_judge(n);
    point_back=modle_judge(n+1);
    if point_front==0 && point_back==0 && point==1
        modle_judge_res(n)=-1;
    end
end
figure
stem(sum_all,"filled","b");
hold on
plot(ones(1,max_num).*threshold_all,"r")
hold on
stem(modle_judge_res.*threshold_all,"filled","y");

ture_infect(find(modle_judge_res==1))=threshold_all;
ture_infect(find(modle_judge_res==-1))=0;
ture_infect(find(ture_infect==0))=NaN;
sum_all(find(sum_all==0))=NaN;
figure
plot(ones(1,max_num-44-10).*threshold_all,"--k")
hold on
bar(1:1:max_num,sum_all,"k");
set(gca,'XTick',1:4:128) 
set(gca,'FontSize',12,'FontName','Arial');
xlabel("Second during contagion in 3 s bins","FontSize",12)%bin
xlim([0,61])
ylabel("Total coughs per bin","FontSize",12)%bin

% hold on
% bar(ture_infect,"k")
% title("Response for cough contagion","FontSize",12,'FontWeight', 'bold')%Infectious judgment
title("Bin=3","FontSize",12,'FontWeight', 'bold')%Infectious judgment
% legend("Threshold","Bin statistic","FontSize",10)%Bin statistic,
txt1 = {'Threshold'};
txt2 = {'Mean = 1.27, SD = 2.77, Threshold = 5.84'};
text(max_num-53,threshold_all,txt1,"FontSize",10)
text(17,max(sum_all)-0.5,txt2,"FontSize",8)
```
