## Supplementary material for "The Evidence of Cough Contagion in Human Beings": Code: account_Boat.m

```
clc;
clear;
close all;
sacle=10;
boat_length1=load("boat_length1.mat","boat_length");
boat_length2=load("boat_length2.mat","boat_length");
boat_length3=load("boat_length3.mat","boat_length");
boat_length4=load("boat_length4.mat","boat_length");
boat_length5=load("boat_length5.mat","boat_length");
boat_length6=load("boat_length6.mat","boat_length");
boat_length7=load("boat_length7.mat","boat_length");
boat_length8=load("boat_length8.mat","boat_length");
boat_length9=load("boat_length9.mat","boat_length");

boat_lengthn1=-sacle*ones(60,1);
boat_lengthn2=-sacle*ones(60,1);
boat_lengthn3=-sacle*ones(60,1);
boat_lengthn4=zeros(20,1);
boat_lengthn5=zeros(20,1);
boat_lengthn6=zeros(20,1);
boat_lengthn7=zeros(20,1);
boat_lengthn1(1:23,1)=boat_length1.boat_length;
boat_lengthn2(1:16,1)=boat_length2.boat_length;
boat_lengthn2(17:26,1)=boat_length3.boat_length;
boat_lengthn3(1:10,1)=boat_length4.boat_length;
boat_lengthn3(11:14,1)=boat_length5.boat_length;
boat_lengthn2(15:30,1)=boat_length6.boat_length;
boat_lengthn2(31:36,1)=boat_length7.boat_length;

boat_lengthn3(15:21,1)=boat_length8.boat_length;
boat_lengthn3(22:34,1)=boat_length9.boat_length;
boat_lengthn=[boat_lengthn1,boat_lengthn2,boat_lengthn3];
figure
name=["Period proportion morning";"Period proportion afternoon"];
for m=2:3
    boat_length=boat_lengthn(:,m);
    boat_length10=round(boat_length/sacle)+1;
    [boat_lengthy,~]=size(boat_length);
    [boat_lengthy10,~]=size(boat_length10);
    array=zeros(uint32(max(boat_length)),1);
    array10=zeros(uint32(max(boat_length10)),1);
    for n=1:boat_lengthy
        if  uint32(boat_length(n))==0
            continue
        end
        array(uint32(boat_length(n)))=array(uint32(boat_length(n)))+1;
        array10(uint32(boat_length10(n)))=array10(uint32(boat_length10(n)))+1;
    end
    array10=array10/(array10'*ones(size(array10)));
    array10(find(array10==0))=NaN;
%     figure
    x=sacle/2:sacle:sacle*max(boat_length10)-sacle/2;
%     [~,x_nx]=size(x);
%     x_t=zeros(1,2*x_nx-1);
%     x_t(1:2:2*x_nx-1)=x+1;
%     x_t(2:2:2*x_nx-2)=x(2:end)-1;
%     x_t=[x_t,x_t(1,end)+8]
%     array10n=zeros(2*max(boat_length10),1);
%     array10n(1:2:2*max(boat_length10)-1,1)=array10;
%     array10n(2:2:2*max(boat_length10),1)=array10;
%     area(x_t,array10n)
%     ylim([0,1])
%     name=strcat("第",num2str(m),"组");
%     title(name);
    subplot (1,2,m-1)
    array10=floor(array10.*100)/100;
    b=bar(x,array10);

    set(gca,'XTick',0:sacle:200) 
    title(name(m-1,1),"FontSize",20);
    xlabel("Boat Duration","FontSize",14)%bin
    ylabel("Probability","FontSize",14)%bin
    set(gca,'FontSize',14,'FontName','Arial');
    ylim([0,1])
    xlim([-10,200])
    xtips1 = b(1).XEndPoints;
    ytips1 = b(1).YEndPoints;
    labels1 = string(b(1).YData);
    text(xtips1,ytips1,labels1,'HorizontalAlignment','center',...
        'VerticalAlignment','bottom')


end
```
