## Supplementary material for "The Evidence of Cough Contagion in Human Beings": Code: aclu.m

```
clc;
clear;
close all;

data=readmatrix("boutXX.xlsx");
data_sex_man=data(find(data(:,4)==1),1);
data_sex_woman=data(find(data(:,4)==2),1);
data_time_am=data(find(data(:,5)==1),1);
data_time_pm=data(find(data(:,5)==0),1);


mean_N_man=mean(data_sex_man);
mean_N_woman=mean(data_sex_woman);
mean_N_am=mean(data_time_am);
mean_N_pm=mean(data_time_pm);

std_N_man=std(data_sex_man);
std_N_woman=std(data_sex_woman);
std_N_am=std(data_time_am);
std_N_pm=std(data_time_pm);


threshold_man=mean_N_man+1.65*std_N_man;
threshold_woman=mean_N_woman+1.65*std_N_woman;
threshold_am=mean_N_am+1.65*std_N_am;
threshold_pm=mean_N_pm+1.65*std_N_pm;
```
