## Supplementary material for "The Evidence of Cough Contagion in Human Beings": Code: Boat.m

```
useful_data_t_type=zeros(useful_datay,2);
for n=1:1:useful_datay
    useful_data_t_type(n,1)=useful_data(n).time;
    useful_data_t_type(n,2)=useful_data(n).type;

end
useful_data_t1=useful_data_t_type(find(useful_data_t_type(:,2)==1));
useful_data_t3=useful_data_t_type(find(useful_data_t_type(:,2)==3));
boat_length=useful_data_t3-useful_data_t1;
 save("D:\XXX晗\XX\boat_length2",'boat_length')
```
