## Supplementary material for "The Evidence of Cough Contagion in Human Beings": Code: definction.m

```
useful_data=data(find(type~=0));

%lonely cough,trigger,cough MPOC,ending cough(Undefined for 0, 1, 2, 3, and 9 respectively)
%The first one to appear is trigger cough
useful_data(1).type=1;
%The last one to appear is ending cough
useful_data(end).type=3;
[useful_datay,~]=size(useful_data);
for n=1:1:useful_datay
    temp_time(1,n)=useful_data(n,1).time;
    temp_number(1,n)=useful_data(n,1).number;
    temp_type(1,n)=useful_data(n,1).type;
end
%judge of trigger cough
for n=2:1:useful_datay-1
    flag_frontm=0;
    flag_fronts=0;
    number_Mutual=temp_number(find(temp_time(1:n-1)>temp_time(n)-Mutual_infection));
    number_Mutual=number_Mutual(find(number_Mutual~=useful_data(n).number));
    number_Self=temp_number(find(temp_time(1:n-1)>temp_time(n)-Self_infection));
    number_Self=number_Self(find(number_Self==useful_data(n).number));
    if isempty(number_Mutual)==1
        flag_frontm=1;
    end
    if isempty(number_Mutual)==0
        flag_frontm=0;
    end

    if isempty(number_Self)==1
        flag_fronts=1;
    end
    if isempty(number_Self)==0
        flag_fronts=0;
    end
    if flag_frontm*flag_fronts==1
        useful_data(n).type=1;
    end
end
%judge of ending cough
for n=2:1:useful_datay-1
    flag_frontm=0;
    flag_fronts=0;
    number_Mutual=temp_number(find(temp_time(n+1:end)<(temp_time(n)+Mutual_infection))+n);
    number_Mutual=number_Mutual(find(number_Mutual~=useful_data(n).number));
    number_Self=temp_number(find(temp_time(n+1:end)<temp_time(n)+Self_infection)+n);
    number_Self=number_Self(find(number_Self==useful_data(n).number));
    if isempty(number_Mutual)==1
        flag_frontm=1;
    end
    if isempty(number_Mutual)==0
        flag_frontm=0;
    end

    if isempty(number_Self)==1
        flag_fronts=1;
    end
    if isempty(number_Self)==0
        flag_fronts=0;
    end
    if flag_frontm*flag_fronts==1
        useful_data(n).type=3;
    end
end
%judge of MPOC
for n=1:1:useful_datay
    if (useful_data(n).type)==9
        useful_data(n).type=2;
    end
end
%judge PSS(1,2,3)
useful_type=zeros(2,useful_datay);
for n=1:1:useful_datay-1
    if useful_data(n).type==2&&useful_data(n+1).type==1
        useful_data(n).type=3;
    end
end
for n=2:1:useful_datay
    if useful_data(n).type==3&&useful_data(n-1).type==3
        useful_data(n-1).type=2;
    end
end
for n=2:1:useful_datay
    if useful_data(n).type==2&&useful_data(n-1).type==3
        useful_data(n-1).type=1;
    end
end
for n=2:1:useful_datay
    if useful_data(n).type==1&&useful_data(n-1).type==1
        useful_data(n).type=2;
    end
end
% useful_data(8).type=2;
for n=1:1:useful_datay
    useful_type(1,n)=useful_data(n).type;
    useful_type(2,n)=useful_data(n).sex;
end
trigger_Loc=0;
MOPC_Loc=0;
ending_Loc=0;

% for useful_num=1:1:useful_datay
%     if useful_type(1,useful_num)==1&&useful_type(2,useful_num)<=3
%         trigger_Loc_temp=useful_num;
% 
%         scanning_on=1;
%         scanning_step=1;
%         while scanning_on
%             if useful_type(1,useful_num+scanning_step)==2
%                 MOPC_Loc=[MOPC_Loc,useful_num+scanning_step];
%             end
%             if useful_type(1,useful_num+scanning_step)==3
%                 ending_Loc=[ending_Loc,useful_num+scanning_step];
%                 scanning_on=0;
%                 break;
%             end
%            scanning_step=1+scanning_step;
%         end
%         trigger_Loc=[trigger_Loc,trigger_Loc_temp];
%     end
% 
% end
% trigger_Loc=trigger_Loc(2:end);
% 
% MOPC_Loc=MOPC_Loc(2:end);
% ending_Loc=ending_Loc(2:end);

trigger_Loc=find(useful_type(1,:)==1);
MOPC_Loc=find(useful_type(1,:)==2);
ending_Loc=find(useful_type(1,:)==3);


% save("useful_data2.mat","useful_data")
```
