## Supplementary material for "The Evidence of Cough Contagion in Human Beings": Code: heat_hist.m

```
clc;
clear;
close all;

A=[1,0,0,0,0,0;...
    -0.040,1,0,0,0,0;...
    0.792,0.339,1,0,0,0;...
    0.747,0.082,0.608,1,0,0;...
    0.538,0.222,0.520,0.716,1,0;...
    -0.645,0.15,-0.479,-0.035,-0.044,1;];
xvalues = {'Number of cough','Response time','Length of chain','Diversity of chain',...
    'Density of chain','Range of chain'};
yvalues = {'Number of cough','Response time','Length of chain','Diversity of chain',...
    'Density of chain','Range of chain'};
h=heatmap(xvalues,yvalues,A);
h.Title = 'Correlation Coefficient heat map';
h.ColorLimits=[-1,1];
h.FontSize=12;


data=readmatrix("bout数据.xlsx");
```
