## Supplementary material for "The Evidence of Cough Contagion in Human Beings": Code: judgeT1_T2.m

```
clc;
clear;
close all;

data_xlex1=readmatrix("D:\XXX\XX\COUGH.xlsx","Sheet","Sheet1");
data_xlex2=readmatrix("D:\XXX\XX\COUGH.xlsx","Sheet","Sheet2");
data_xlex3=readmatrix("D:\XXX\XX\COUGH.xlsx","Sheet","Sheet3");
data_xlex4=readmatrix("D:\XXX\XX\COUGH.xlsx","Sheet","Sheet4");
data_xlex5=readmatrix("D:\XXX\XX\COUGH.xlsx","Sheet","Sheet5");
data_xlex6=readmatrix("D:\XXX\XX\COUGH.xlsx","Sheet","Sheet6");
data_xlex7=readmatrix("D:\XXX\XX\COUGH.xlsx","Sheet","Sheet7");

[data_xlex1_y,~]=size(data_xlex1);
[data_xlex2_y,~]=size(data_xlex2);
[data_xlex3_y,~]=size(data_xlex3);
[data_xlex4_y,~]=size(data_xlex4);
[data_xlex5_y,~]=size(data_xlex5);
[data_xlex6_y,~]=size(data_xlex6);
[data_xlex7_y,~]=size(data_xlex7);
```
