## Supplementary material for "The Evidence of Cough Contagion in Human Beings": Code: model_function.m

```
clc
clear;
close all;
Mutual_infection=30;%Time of mutual infection (s)
Self_infection=45; %Self transmission time (s)
sum_N_mat=zeros(7,200);
bin=3;

namenum=7;
Sheetnamenum=strcat("Sheet",num2str(namenum));

    data_xlex=readmatrix("D:\XXX\XX\COUGH.xlsx","Sheet",Sheetnamenum);
    [time_data,xlabel_data]=size(data_xlex);
    time=data_xlex(:,3);
    for n=1:1:time_data
        data(n,1)=struct('time',data_xlex(n,3),'number',data_xlex(n,2),'type',0,'sex',data_xlex(n,4),'am_pm',data_xlex(n,5));
    end
    number=data_xlex(:,2);
    %pretreatment
    %lonely cough,trigger,cough MPOC,ending cough(Undefined for 0, 1, 2, 3, and 9 respectively)
        for n=1:1:time_data
            if n==1
                timefront=data(n);
                timebegin=data(n);
                timeback=data(n+1);
                Self_expess_lim=timebegin.time+Self_infection;%Autobroadcast critical time (relative to start time)
                Mutual_expess_lim=timebegin.time+Mutual_infection;%Cross propagation critical time (relative starting time)
                [point]=time(time<Mutual_expess_lim);%All data before the critical time of mutual propagation
                line_self_expess=number(find(time(n+1:end)<time(n)+Self_infection)+n);
                line_Mutual_expess=number(find(time(n+1:end)<time(n)+Mutual_infection)+n);%Find all events that are less than the mutual propagation time
                event=data(n).number;
                line_Mutual_expess=line_Mutual_expess(find(line_Mutual_expess~=event));
                line_self_expess=line_self_expess(find(line_self_expess==event));
                returnValueS = isempty(line_self_expess);%The judgment matrix is empty
                if returnValueS==1
                    flag_M=1;
                end
                if returnValueS==0
                    flag_M=0;
                end
                returnValueM = isempty(line_Mutual_expess);%The judgment matrix is empty
                if returnValueM==1
                    flag_S=1;
                end
                if returnValueM==0
                    flag_S=0;
                end
                if flag_M*flag_S==1
                    data(n).type=0;
                end
                if flag_M*flag_S==0
                    data(n).type=9;
                end
                type=data(n).type;
            end
            if n~=1
                Self_expess_lim=timebegin.time+Self_infection;%Autobroadcast critical time
                Mutual_expess_lim=timebegin.time+Mutual_infection;%Cross propagation critical time
                time_limS_front=time(n)-Self_infection;
                time_limS_back=time(n)+Self_infection;
                time_limM_front=time(n)-Mutual_infection;
                time_limM_back=time(n)+Mutual_infection;
                time_BigSelf_1=time(find(time~=time(n)));
                time_BigSelf_1=time_BigSelf_1(find(time_BigSelf_1>time_limS_front));
                line_self_expess=number(find(time_BigSelf_1<time_limS_back));
                time_BigMutual=time(find(time~=time(n)));
                time_BigMutual=time_BigMutual(find(time_BigMutual>time_limM_front));
                line_Mutual_expess=number(find(time_BigMutual<time_limM_back));
                event=data(n).number;
                line_Mutual_expess=line_Mutual_expess(find(line_Mutual_expess~=event));
                line_self_expess=line_self_expess(find(line_self_expess==event));
                returnValueS = isempty(line_self_expess);%The judgment matrix is empty
                if returnValueS==1
                    flag_M=1;
                end
                if returnValueS==0
                    flag_M=0;
                end
                returnValueM = isempty(line_Mutual_expess);%The judgment matrix is empty
                if returnValueM==1
                    flag_S=1;
                end
                if returnValueM==0
                    flag_S=0;
                end
                if flag_M*flag_S==1
                    data(n).type=0;
                end
                if flag_M*flag_S==0
                    data(n).type=9;
                end
                type=[type;data(n).type];
            end
    
        end

definction
Statistics
```
