## Supplementary material for "The Evidence of Cough Contagion in Human Beings": Code: num_length_peruid_sex.m

```
clc;
clear;
close all;
useful_data2=load("useful_data2.mat","useful_data");
useful_data3=load("useful_data3.mat","useful_data");
useful_data4=load("useful_data4.mat","useful_data");
useful_data5=load("useful_data5.mat","useful_data");
useful_data6=load("useful_data6.mat","useful_data");
useful_data7=load("useful_data7.mat","useful_data");
useful_data8=load("useful_data8.mat","useful_data");
useful_data9=load("useful_data9.mat","useful_data");

useful_data2=useful_data2.useful_data;
useful_data3=useful_data3.useful_data;
useful_data4=useful_data4.useful_data;
useful_data5=useful_data5.useful_data;
useful_data6=useful_data6.useful_data;
useful_data7=useful_data7.useful_data;
useful_data8=useful_data8.useful_data;
useful_data9=useful_data9.useful_data;

useful_data=[useful_data2;useful_data3;useful_data4;useful_data5;useful_data6;...
    useful_data7;useful_data8;useful_data9];
[data_y,data_x]=size(useful_data);
bout=zeros(82,5);
m=0;
for n=1:data_y
    if useful_data(n).type==1
        time_begin=useful_data(n).time;
        time_MPOC=useful_data(n+1).time;
        trigger_sex=useful_data(n).sex+1;
        m=m+1;
        bout(m,1)=bout(m,1)+1;%Calculate the number of coughs
        bout(m,3)=time_MPOC-time_begin;%Calculate the start time of infection
        bout(m,4)=trigger_sex;
        bout(m,5)=useful_data(n).am_pm;
    end
    if useful_data(n).type==2
        bout(m,1)=bout(m,1)+1;%Calculate the number of coughs
    end
    if useful_data(n).type==3
        bout(m,1)=bout(m,1)+1;%Calculate the number of coughs
        time_end=useful_data(n).time;
        bout(m,2)=time_end-time_begin;%Calculate the duration of bouts
    end
end


writematrix(bout,"D:\XXX\XX\boutXX.xlsx" ,'Sheet',1,'Range','A2:E83');
```
