## Supplementary material for "The Evidence of Cough Contagion in Human Beings": Code: pin.m

```
clc;
clear;
close all;

useful_data2=load("useful_data2.mat","useful_data");
useful_data3=load("useful_data3.mat","useful_data");
useful_data4=load("useful_data4.mat","useful_data");
useful_data5=load("useful_data5.mat","useful_data");
useful_data6=load("useful_data6.mat","useful_data");
useful_data7=load("useful_data7.mat","useful_data");
useful_data8=load("useful_data8.mat","useful_data");
useful_data9=load("useful_data9.mat","useful_data");

useful_data2=useful_data2.useful_data;
useful_data3=useful_data3.useful_data;
useful_data4=useful_data4.useful_data;
useful_data5=useful_data5.useful_data;
useful_data6=useful_data6.useful_data;
useful_data7=useful_data7.useful_data;
useful_data8=useful_data8.useful_data;
useful_data9=useful_data9.useful_data;


[x2,y2]=size(useful_data2);
[x3,y3]=size(useful_data3);
[x4,y4]=size(useful_data4);
[x5,y5]=size(useful_data5);
[x6,y6]=size(useful_data6);
[x7,y7]=size(useful_data7);
[x8,y8]=size(useful_data8);
[x9,y9]=size(useful_data9);
for n=1:x2
    data2(n,1)=useful_data2(n).type;
end
for n=1:x3
    data3(n,1)=useful_data3(n).type;
end
for n=1:x4
    data4(n,1)=useful_data4(n).type;
end
for n=1:x5
    data5(n,1)=useful_data5(n).type;
end
for n=1:x6
    data6(n,1)=useful_data6(n).type;
end
for n=1:x7
    data7(n,1)=useful_data7(n).type;
end
for n=1:x8
    data8(n,1)=useful_data8(n).type;
end
for n=1:x9
    data9(n,1)=useful_data9(n).type;
end

num2=zeros(size(find(data2==1)));
num3=zeros(size(find(data3==1)));
num4=zeros(size(find(data4==1)));
num5=zeros(size(find(data5==1)));
num6=zeros(size(find(data6==1)));
num7=zeros(size(find(data7==1)));
num8=zeros(size(find(data8==1)));
num9=zeros(size(find(data9==1)));
num_all=size(find(data2==1))+size(find(data3==1))+size(find(data4==1))+size(find(data5==1))+size(find(data6==1))+size(find(data7==1))+size(find(data8==1))+size(find(data9==1));
m=0;
for n=1:size(data2)
    if data2(n)==1
        m=m+1;
        num2(m)=num2(m)+1;
    end
    if data2(n)~=1
        num2(m)=num2(m)+1;
    end
end
m=0;
for n=1:size(data3)
    if data3(n)==1
        m=m+1;
        num3(m)=num3(m)+1;
    end
    if data3(n)~=1
        num3(m)=num3(m)+1;
    end
end
m=0;
for n=1:size(data4)
    if data4(n)==1
        m=m+1;
        num4(m)=num4(m)+1;
    end
    if data4(n)~=1
        num4(m)=num4(m)+1;
    end
end
m=0;
for n=1:size(data5)
    if data5(n)==1
        m=m+1;
        num5(m)=num5(m)+1;
    end
    if data5(n)~=1
        num5(m)=num5(m)+1;
    end
end
m=0;
for n=1:size(data6)
    if data6(n)==1
        m=m+1;
        num6(m)=num6(m)+1;
    end
    if data6(n)~=1
        num6(m)=num6(m)+1;
    end
end
m=0;
for n=1:size(data7)
    if data7(n)==1
        m=m+1;
        num7(m)=num7(m)+1;
    end
    if data7(n)~=1
        num7(m)=num7(m)+1;
    end
end
m=0;
for n=1:size(data8)
    if data8(n)==1
        m=m+1;
        num8(m)=num8(m)+1;
    end
    if data8(n)~=1
        num8(m)=num8(m)+1;
    end
end
m=0;
for n=1:size(data9)
    if data9(n)==1
        m=m+1;
        num9(m)=num9(m)+1;
    end
    if data9(n)~=1
        num9(m)=num9(m)+1;
    end
end
num_all=[num2;num3;num4;num5;num6;num7;num8;num9];
all=num_all'*ones(82,1);
%Calculate mean
mean_N=mean(num_all);
%Calculate standard deviation
std_N=std(num_all);
```
