## Supplementary material for "The Evidence of Cough Contagion in Human Beings": Code: Statistics.m

```
% bin=3;
% bin=5;
[~,bout_y]=size(trigger_Loc);
bout=zeros(2,bout_y);
bout(1,:)=temp_time(trigger_Loc);
bout(2,:)=temp_time(ending_Loc);
period=bout(2,:)-bout(1,:);
N_size=ceil(period/bin);
N_size_max=max(N_size);

bout_number_list=zeros(bout_y,N_size_max);
bout_list=zeros(1,ceil(temp_time(1,end)/bin+1)*bin);
bout_list(uint32(temp_time(trigger_Loc))+1)=1;
bout_list(uint32(temp_time(MOPC_Loc))+1)=2;
bout_list(uint32(temp_time(ending_Loc))+1)=3;
% plot(bout_list)
for n=1:1:bout_y %统计每一个bout内bin值0
    for m=1:1:N_size(n)%统计bin内MOPC与ending cough数量
        [~,num_MPOC]=size(find(bout_list(round(bout(1,n))+(m-1)*bin+1:round(bout(1,n))+(m)*bin)==2));
        [~,num_ending]=size(find(bout_list(round(bout(1,n))+(m-1)*bin+1:round(bout(1,n))+(m)*bin)==3));
        bout_number_list(n,m)=num_MPOC+num_ending;
    end
end
%bout相同时间点的bin中的统计量加和在一起
accum=ones(bout_y,1);
sum_N=(bout_number_list'*accum)';
%计算均值
mean_N=mean(sum_N);
%计算标准差
std_N=std(sum_N);
%阈值
threshold=mean_N+1.65*std_N;
ture_infect=zeros(1,N_size_max);
ture_infect(find(sum_N>=threshold))=sum_N(find(sum_N>=threshold));
%不显示0值
sum_N(find(sum_N)==0)=NaN;
ture_infect(find(ture_infect)==0)=NaN;


% figure
% stem(sum_N,"filled","b");
% hold on
% plot(ones(1,N_size_max).*threshold,"r")
% hold on
% stem(ture_infect,"g",'MarkerSize',10,'LineWidth',1)
% title("传染性判断")
% legend("bin统计量","阈值","传染性")
savestrss=["sum_N_4.14_1.mat";...
                "sum_N_4.14_2.mat";...
                "sum_N_4.18_1.mat";...
                "sum_N_4.18_2.mat";...
                "sum_N_4.23_1.mat";...
                "sum_N_4.23_2.mat";...
                "sum_N_4.25_1.mat";...
                "sum_N_4.25_2.mat"];
save(savestrss(namenum-1),'sum_N');
```
